# Effects of species invasion on population dynamics, vital rates, and life histories of the native species

**DOI:** 10.1101/177741

**Authors:** Simone Vincenzi, Alain J Crivelli, Dusan Jesensek, Ellen Campbell, John C Garza

## Abstract

Invasions occurring in natural environments provide the opportunity to study how vital rates change and life histories evolve in the presence of a competing species. In this work, we estimate differences in reproductive traits, individual growth trajectories, survival, life histories, and population dynamics between a native species living in allopatry and in sympatry with an invasive species of the same taxonomic Family. We used as a model system marble trout *Salmo marmoratus* (native species) and rainbow trout *Oncorhynchus mykiss* (non-native) living in the Idrijca River (Slovenia). An impassable waterfall separates the stream in two sectors only a few hundred meters apart: a downstream sector in which marble trout live in sympatry with rainbow trout and a upstream sector in which marble trout live in allopatry. We used an overarching modeling approach that uses tag-recapture and genetic data (> 2,500 unique marble and rainbow trout were sampled and SNP-genotyped) to reconstruct pedigrees, test for synchrony of population dynamics, and model survival and growth while accounting for individual heterogeneity in performance. The population dynamics of the two marble trout populations and of rainbow trout were overall synchronous. We found higher prevalence of younger parents, higher mortality, and lower population density in marble trout living in sympatry with rainbow trout than in marble trout living in allopatry. There were no differences in the average individual growth trajectories between the two marble trout populations. Faster life histories of marble trout living in sympatry with rainbow trout are consistent with predictions of life-history theory.

## Introduction

The invasion of native communities by alien species is one of the major threats to the stability of ecosystem processes and the persistence of endemic species (Simon and Townsend 2003). Although they are one of the biggest challenges conservation biologists are facing (Allendorf and Lundquist 2003), invasions occurring in natural environments also provide the opportunity to study how vital rates change and life histories (the timing of key events in an organism’s lifetime) evolve in competing species, a central topic of ecology and evolutionary biology (Sakai et al. 2001). In particular in the case of long-lived and mobile organisms, insights from laboratory or small-scale experiments are of limited value, since spatial constraints may force species to interact at unnaturally small scales (Korsu et al. 2009), the effects of the invasion on the native communities may be appreciated only after a few generations, and the size and direction of the effects may change with changes in the environment (Carroll 2007).

The study of biological invasions has mostly focused on understanding the mechanism by which non-native species successfully invade new regions. In particular, the focus has been on the environmental (e.g. climate, Jiménez *et al.* 2011) or ecological (e.g. unoccupied niches, Olden *et al.* 2006) conditions facilitating or hampering establishment and spread of the invading species, the life-history traits that are associated with invasion success (Capellini et al. 2015), and the adoption of life-history strategies by the colonist population that differ from those of the source population (Amundsen et al. 2012). On the other hand, few studies have investigated the overall effects of invasion on the native species, such as effects on its diet, growth, survival, reproduction, life histories, and population dynamics (Peterson et al. 2004).

In this work, we combine a fine-grained, long-term tag-recapture dataset with statistical and modeling methods that take into account individual and temporal heterogeneity in vital rates to estimate differences in individual growth, survival, reproduction, life histories, and population dynamics between individuals of a native species living in allopatry or competing with an invasive species, using as model system marble trout *Salmo marmoratus* (native species) and rainbow trout *Oncorhynchus mykiss* (non-native species) living in Western Slovenian streams. Salmonids have been often used as model systems in ecology and evolutionary biology, due to their vast geographic distribution, their ecological and life-history variability, and their strong genetic and plastic responses to habitat and climate variation (Elliott 1994; Stearns and Hendry 2003; Garcia de Leaniz et al. 2007; Jonsson and Jonsson 2011).

Marble trout is endemic to rivers tributary to the upper Adriatic Sea and persists in pure form in the Alps of Western Slovenia and parts of Northern Italy (Sušnik et al. 2015; Vincenzi et al. 2016b). Two of the eight remaining genetically pure populations of marble trout in Western Slovenia are in the Idrijca River, where an impassable waterfall separates fish in Upper Idrijca from a closely related group in Lower Idrijca (Fumagalli et al. 2002). Marble trout in Lower Idrijca co-exist with non-native rainbow trout, which were introduced in the 1960s and have been established since then, but they are absent in Upper Idrijca (Stanković et al. 2015). Given the high similarity of Lower and Upper Idrijca habitats (Vincenzi et al. 2016b), this unintended treatment-control experiment allows studying variation in vital rates, life histories, and population dynamics of the native species when competing with an invasive species of the same taxonomic Family.

The absence of replicates does not allow for strong inference (*sensu* Platt 1964) on the causes of variation in vital rates, life histories, and population dynamics, therefore we use the result of the study not to test, but to generate hypotheses on the ultimate and proximate causes of the observed variation. Specifically, we used tag-recapture and genetic data (> 2,500 unique marble and rainbow trout were sampled and SNP-genotyped) to reconstruct pedigrees and estimate age of parents, model survival using tag-recapture analysis, and model growth using random-effects models that take explicitly into account individual heterogeneity in performance. The overarching data and modeling approach allowed us to test for population synchrony, and estimate differences in population dynamics, proportion of younger parents, individual growth trajectories, and survival between marble trout living in allopatry and in sympatry with rainbow trout. We then discuss the observed variation in life histories in the light of ecological and life-history theory.

## Material and Methods

### Study area and species description

The Idrijca catchment is pristine and there are neither poaching nor angling in the stream. Marble trout is the only fish species living in Upper Idrijca. Within Lower (LIdri) and Upper (UIdri) Idrijca, there are no physical barriers impairing upstream or downstream movement of marble and rainbow trout; however, both species are territorial and their movement throughout their lifetime is typically limited (Vincenzi et al. 2016b). Mean daily water temperature in LIdri and UIdri over years 2006-2013 were 8.1±0.2 and 7.7±0.2 °C (Vincenzi et al. 2016b). There are ample spawning grounds in both Lower and Upper Idrijca.

Detailed description of the biology and ecology of marble trout can be found in (Vincenzi et al. 2016b). Marble trout populations are highly genetically differentiated (Fumagalli et al. 2002), persist at low population densities (between ∼600 and 1250 fish ha^-1^ for fish older than young-of-year), are at high risk of extinction due to hybridization with non-native brown trout (*S. trutta*), and the occurrence of extreme events such as flash floods, debris flows, and landslides (Vincenzi et al. 2008; Vincenzi et al. 2016b; Vincenzi et al. 2017).

Rainbow trout is a north Eastern Pacific species (Gall and Crandell 1992) that was introduced in the Adriatic basin of Slovenia in the early 20^th^ century and there established self-sustaining stream-resident populations (Stanković et al. 2015). Rainbow trout was stocked in the headwaters of the Idrijca River (Upper Idrijca) only once, in 1962. Rainbow trout in the Adriatic basin of Slovenia typically start spawning at 1 year old (D. Jesensek & A.J. Crivelli, *unpublished data*).

#### Sampling

We sampled the rainbow and marble populations of LIdri and UIdri bi-annually in June and September of each year from June 2004 to September 2015 for a total of 24 sampling occasions. Fish were captured by electrofishing and total length (*L*) and weight (*W*) recorded to the nearest mm and g. If captured fish had *L* > 115 mm, and had not been previously tagged or had lost a previously applied tag, they received a Carlin tag (Carlin 1955) and age was determined by reading scales. Fish are aged as 0+ in the first calendar year of life, 1+ in the second year and so on. Sub-yearlings of both rainbow and marble trout are smaller than 115 mm in June and September, so fish were tagged when at least aged 1+. The adipose fin was also removed from all fish captured for the first time (starting at age 0+ in September), including those not tagged due to small size at age 1+. Therefore, fish with intact adipose fin were not sampled at previous sampling occasions at age 0+ or 1+. Males and females of marble and rainbow trout are morphologically indistinguishable in either June or September; sex was thus assigned in the lab using molecular techniques (Yano et al. 2013).

#### SNP discovery and genotyping

DNA was extracted from dried fin clips using the Dneasy 96 filter-based nucleic acid extraction system on a BioRobot 3000 (Qiagen, Inc.), following the manufacturer’s protocols. Extracted DNA was diluted 2:1 with distilled water and used for polymerase chain reaction (PCR) amplification of 94 population-specific SNPs. SNPs were assayed with 96.96 genotyping IFC chips on an EP1 (EndPoint Reader 1) instrument (Fluidigm, Inc.), using the manufacturer’s recommended protocols. Genotypes were called using SNP Genotyping Analysis software (Fluidigm). Two people called all genotypes independently, and discrepancies in the scores were resolved either by consensus, by re-genotyping, or by deletion of that genotype from that data set. A proportion of individuals were genotyped more than once (e.g. in case of tag loss or when the fish was < 115 mm, and later at tagging), as determined by observed identical genetic profiles and compatible age and length data, and one of the samples was excluded from the analyses.

Matching genotypes of individuals with different tags is a method of “genetic tagging” that allows reconstructing the life histories of individuals after tag loss. Mean±sd minor allele frequency (MAF) of the SNPs was 0.27±0.13 for LIdri_MT (95 polymorphic SNPs, Table EMS1), 0.27±0.14 for UIdri_MT (95 SNPs, Table EMS2), and 0.26±0.14 for LIdri_RT (83 SNPs, Table EMS3). Such MAFs with nearly 100 SNPs are sufficient for parent pair/offspring trio reconstruction with high accuracy (Anderson and Garza 2006). The final dataset included 1129 marble trout living in LIdri (LIdri_MT), 610 marble trout living in UIdri (UIdri_MT), and 291 rainbow trout living in LIdri (LIdri_RT) that were identified either as offspring or potential parents.

### Statistical analyses and hypotheses tested

We first estimated density of age-classes and then tested for (a) recruitment-driven population dynamics, (b) population synchrony, and (c) temporal differences in population densities between the three populations. Then, we reconstructed pedigrees in each population and fitted models of growth and survival to estimate differences in (d) lifetime growth trajectories, (e) probability of survival, and (f) proportion of young parents between LIdri_MT, LIdri_RT, and UIdri_MT.

#### Estimation of population density

We estimated density of 0+ fish only in September, since marble trout fish normally emerge a few days before the June sampling and rainbow trout emerge in July. We estimated density of 0+ fish and fish older than 0+ using a two-pass removal protocol (Carle and Strub 1978) as implemented in the R (R Development Core Team 2014) package FSA (Ogle 2015). Total stream surface of the monitored area (1084 m^2^ and 1663 m^2^ for LIdri and UIdri, respectively) was used for the estimation of fish density (in fish ha^-1^). Following Vincenzi et al. (2016b), we tested for recruitment-driven population dynamics in LIdri_MT, LIdri_RT, UIdri_MT by estimating cross-correlations between density of 0+ fish (*D*_0+_) in September (recruitment) and density of older than 0+ (*D*_>0+_) one year later within each population. We also tested for synchrony of population dynamics (“Moran effect”, Ranta *et al.* 1997) of LIdri_MT and LIdri_RT, and LIdri_MT and UIdri_MT. Statistical significance was set at the 0.05 level.

#### Pedigree reconstruction and age at spawning

We used a two-step, conservative approach for reconstructing pedigrees in the marble and rainbow trout populations (Vincenzi et al. 2017), using the software SNPPIT (Anderson 2012) and FRANz (Riester et al. 2009). SNPPIT is tailored for inference of mother-father parent pairs in single generations and does not perform single-parent assignments. FRANz can also infer single parents and can be used to infer parentage in multiple generations. Known parent-offspring relationships can be used by FRANz to estimate genotyping error rate. Marble trout genotypes were analyzed with SNPPIT to identify parent-pair/offspring trios with a false discovery rate threshold of 0.05, which were then used in FRANz as known relationships to identify single parent-offspring pairs and additional parent-pair-offspring trios. Further details on pedigree reconstruction are in Text EMS1.

Marble trout in UIdri and LIdri spawn from mid-November to mid-December and offspring emerge in June. Therefore, say, the eggs of cohort 2011 were fertilized in November-December of 2010 and newborn emerged in June 2011. Rainbow trout in Lower Idrijca spawn in April and offspring emerge in July of the same year (T.W. Lee & A.J. Crivelli, *unpublished data*). Marble trout are iteroparous. Age at first reproduction varies within and among marble trout populations; pedigree reconstruction in other populations found that marble trout spawn typically at age 3+ and older, with fewer fish spawning at age 1+ and 2+ (Meldgaard et al. 2007; Vincenzi et al. 2017).

We used a χ^2^ test of equality of proportions to test for differences in the proportion of younger parents between LIdri_MT and UIdri_MT. Since the distribution of age at spawning is skewed and potentially biased by some old parents, we only considered parents aged 1+ and 2+ (young parents), and 3+ and 4+ (old parents).

#### Growth and body size

To characterize size-at-age and growth trajectories, we modeled variation in lifetime growth trajectories of marble and rainbow trout living in LIdri_MT, LIdri_RT, and UIdri_MT taking into account within-population individual heterogeneity in growth.

The standard von Bertalanffy model for growth (vBGF von Bertalanffy 1957) is

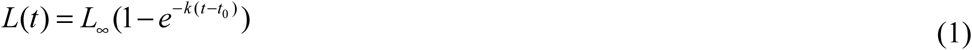

where *L*_∞_ is the asymptotic size, *k* is a coefficient of growth (in time^-1^), and *t*_0_ is the (hypothetical) age at which length is equal to 0. However, to accurately estimate overall, group-specific, or individual growth trajectories we need to take into account individual variation in growth, otherwise the average trajectory can be substantially pulled up or down by a few long-living fast or slow growers (Vincenzi et al. 2014).

We used the formulation of the vBGF specific for longitudinal data of Vincenzi et al. (Vincenzi et al. 2014), in which *L*_∞_ and *k* may be allowed to be a function of shared predictors and individual random effects. In the estimation procedure, we used a log-link function for *k* and *L*_∞_, since both parameters must be non-negative. We set

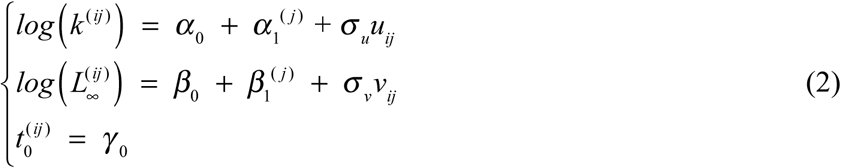

where *u* ∼ *N* (0,1) and *v* ∼ *N* (0,1) are the standardized individual random effects, 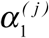 and 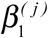 are group effects (e.g. sex, year-of-birth cohort, population, species),*σu* and *σ v* are the standard deviations of the statistical distributions of the random effects, and the other parameters are defined as in Eq. (1).

We thus assume that the observed length of individual *i* in group *j* at age *t* is

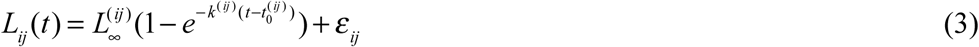

where *ε_ij_* is normally distributed with mean 0 and variance 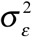.

Since the growth model operates on an annual time scale and more data on tagged fish were generally available in September of each year, we used September data for modeling lifetime growth trajectories. In order to test for differences in growth trajectories, we included three potential categorical predictors (*α*_1_ and β_1_ in Eq. 2) of *k* and *L*_∞_ : (*i*) *Species* (MT and RT), (*ii*) *Population* (Lidri_MT, UIdri_MT, LIdri_RT), (*iii*) *Cohort*. We also tested whether there were large differences in vBGF population-specific models when estimating parameters separately for each population using a standard non-linear regression routine with no random effects (*nls* function in R) or when using models in Eqs. (2) and (3).

#### Survival

Our goal was to estimate variation in survival among populations and to investigate the effects on survival of presence of rainbow trout, time (sampling occasion or season), and successful reproduction.

Two probabilities can be estimated from a capture history matrix: ϕ, the probability of apparent survival (defined “apparent” as it includes permanent emigration from the study area), and *p*, the probability that an individual is captured when alive (Thomson et al. 2009). We used the Cormack–Jolly–Seber (CJS) model as a starting point for the analyses (Thomson et al. 2009). We tested the goodness-of-fit of the CJS model with the program Release. The global model (i.e. the model with the maximum parameterization) was a good starting point to model survival and capture probabilities. All other survival models tested were simplified versions of this global starting model, with the potential addition of covariates. From the global model, recapture probability was modeled first. The recapture model with the lowest AIC was then used to model survival probabilities.

We modeled the seasonal effect (*Season*) as a simplification of full time variation, by dividing the year into two periods: June to September (*Summer*), and the time period between September and June (*Winter*). As length of the two intervals (*Summer* and *Winter*) was different (3 months and 9 months), we estimated probability of survival on a common annual scale. Previous work has found no or minor effects of population density, water temperature, body size, or sex on survival in marble trout (Vincenzi et al. 2016b).

Since only trout with *L* > 115 mm (aged at least 1+) were tagged, capture histories were generated only for those fish. We first used data for all three populations to estimate the overall differences in survival probabilities among them, using population (LIdri_MT, LIdri_RT, UIdri_MT) and sampling occasion or season as predictors of survival. Then, in order to obtain a finer-grained estimation of variation in survival in marble trout, we excluded LIdri_RT from the dataset and modeled survival only in LIdri_MT and UIdri_MT, also in this case using population (LIdri_MT, UIdri_MT) and sampling occasion or season as predictors of survival.

Next, we used a subset of the tag-recapture data set (fish born after the start of tagging, i.e. >=2004, and only females) for the two marble trout populations to test for costs of reproduction. Spawning successfully (i.e. at least one offspring assigned to the parent) was modeled as a time-varying predictor (*Spawn_temp*: 1 for the September to June sampling period in which spawning occurred and 0 otherwise). The hypothesis tested is of a lower survival probability of female parents in the months following reproduction in LIdri_MT than in UIdri_MT due to higher costs of reproduction. Models tested included a population component (constant or LIdri_MT/UIdri_MT), a time component (sampling occasion or season) and a spawning component (constant or *Spawn_temp)*.

For each model, we tested additive and multiplicative interactions among predictors. Models were considered to provide the same fit to data when ΔAIC between model was < 2 (Burnham and Anderson 2002). We carried out the analysis of survival using the package *marked* (Laake et al. 2013) for R (R Development Core Team 2014).

## Results

A paired *t*-test found significant higher population density in September of marble trout older than 0+ in UIdri_MT and LIdri_MT (*p* = 0.01, average difference ∼250 ind. ha^-1^) and significant higher density of 0+ in LIdri_MT (*p* < 0.01, average difference ∼500 ind. ha^-1^) (Fig. 1a). LIdri_RT older than 0 + were found at much lower densities than LIdri_MT (p< 0.01, average difference ∼550 ind. ha^-1^), but densities of 0+ were not significantly different between LIdri_RT and LIdri_MT (*p* = 0.06) (Fig. 1b).

**Fig. 1.**
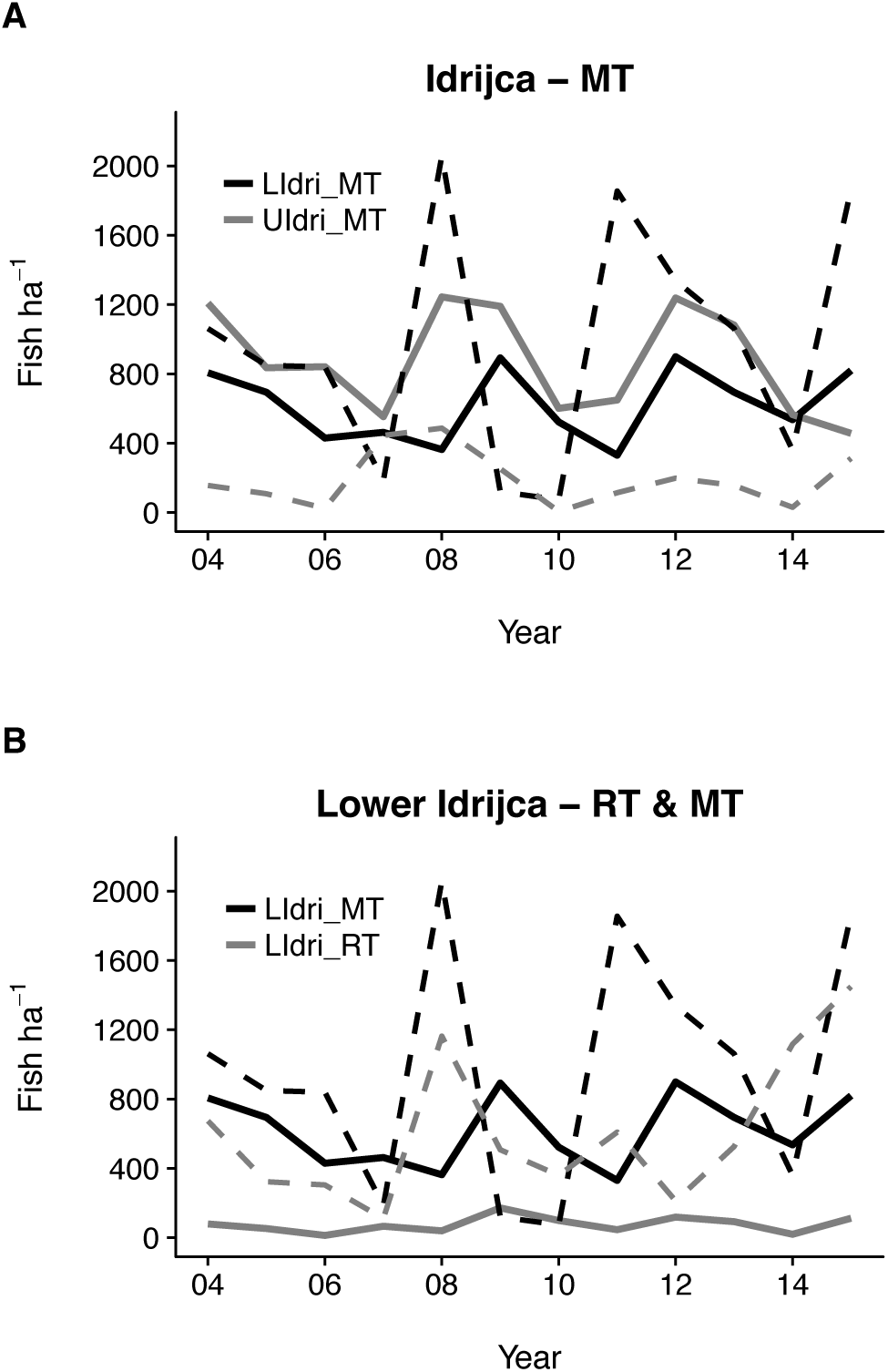
(A) Density over time of LIdri_MT and UIdri_MT aged 0+ (dashed line) and older than 0+ (solid line) in September. (B) Density over time of LIdri_MT and LIdri_RT aged 0+ (dashed line) and older than 0+ (solid line) in September.

There were significant cross-correlations with lag of 1 year between density of 0+ and fish older than 0+ for both LIdri_MT (*r* = 0.65) and UIdri_MT (*r* = 0.61), which suggest that recruitment (i.e. density of 0+ in September at year *t*) was driving variation in density of fish older than juveniles. On the contrary, the one-year lagged correlation between density of 0+ and fish older than 0+ for LIdri_RT was not significant. Densities of LIdri_MT and LIdri_RT older than 0+ were cross-correlated with no lag (*r* = 0.77), while their densities of 0+ were cross-correlated with no lag only when excluding an outlier (year 2012, *r* = 0.60). The densities of both 0+ and older than 0+ of the two marble trout populations (LIdri_MT and UIdri_MT) were not significantly cross-correlated; however, when excluding the outliers 2007 (for 0+) and 2008 (for older than 0+) (which identify the same process, since the 2007 high recruitment in UIdri_MT led to the high density of fish older than 0+ one year later, Fig. 1), the two time series were significantly correlated with no lag (0+: *r* = 0.63, > 0+: *r* = 0.62).

Some old, big marble trout were found in LIdri_MT (Fig. 2), probably due to a large pool in LIdri that allowed their survival. vBGF models fitted with no random effects show how the estimated average growth trajectories can be heavily biased by a few long-living, bigger-at-age marble trout living in Upper and Lower Idrijca (Fig. EMS1), thus making necessary the adoption of models with individual random effects. Random-effects models with *Species* and *Population* as predictors of vBGF’s parameters provided the same fit to data (ΔAIC < 2). Both average growth trajectories and confidence intervals of LIdri_MT and UIdri_MT almost completely overlapped, while on average rainbow trout grew faster and remained bigger-at-age throughout rainbow trout lifetime than marble trout in either population (Fig 2).

**Fig. 2.**
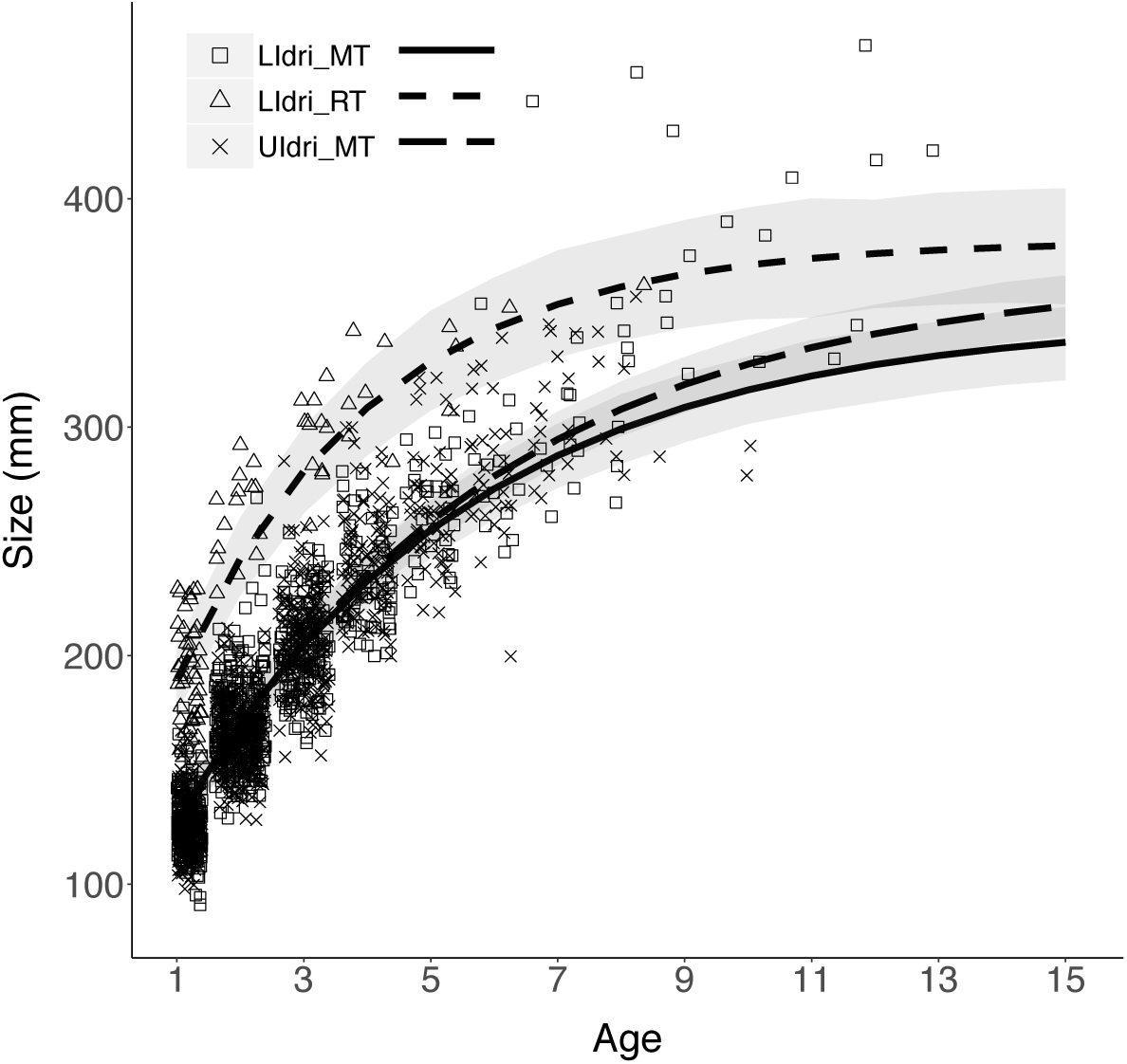
Average (i.e. individual random effects for *L*_∞_ and *k* set to 0) growth trajectories of LIdri_MT, LIdri_RT, and UIdri_MT predicted by the vBGF model with Population as predictor of *L*_∞_ and *k*. Grey bands are the 95% confidence intervals of the average trajectories.

The best model of survival probabilities had survival varying with sampling occasion and population, with survival probabilities consistently higher in UIdri_MT than in LIdri_MT and LIdri_RT (Table 1, Fig. 3a). When including only marble trout populations, the best models had survival varying with population and sampling occasion (Table 1). The addition of *Spawn_temp* did not improve model fit after accounting for population and sampling occasion (Table 1, Fig. 3b).

**Fig. 3.**
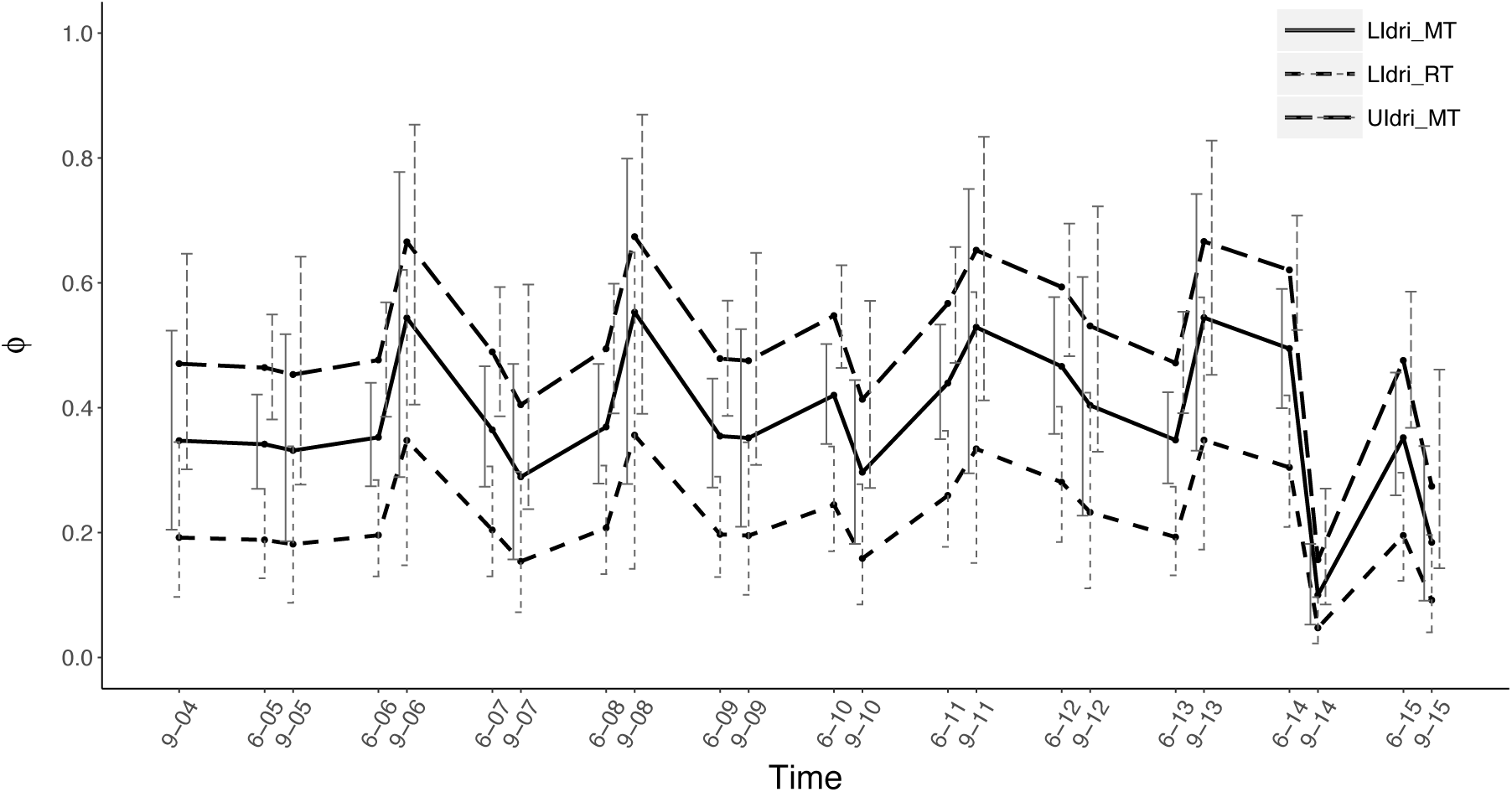
Best models of apparent survival when including all three populations, with survival varying by population and sampling occasion (i.e., *ϕ*(*Population + Time*)).

**Table 1.** Five best models of probability of apparent survival *ϕ* for all three populations (LIdri_MT, UIdri_MT, LIdri_RT), only the two marble trout populations, and only females for the two marble trout populations, using the best model for probability of capture (i.e. *p*(*Pop*)). The symbol * denotes interaction between predictors. *Time* = interval between two consecutive sampling occasions. *Season* = categorical variable for *Summer* (June to September) and *Winter* (September to June). *Spawn_temp* = categorical variable for females spawning successfully (1) or not (0). *npar* = number of parameters of the survival model.

| Model | npar | AIC | ΔAIC | weight |
| --- | --- | --- | --- | --- |
| All three populations |  |  |  |  |
| $\phi (Pop + Time)$ | 28 | 6814.12 | 0.00 | 0.97 |
| $\phi (Pop * Season)$ | 9 | 6820.99 | 6.87 | 0.03 |
| $\phi (Pop)$ | 6 | 6827.94 | 13.82 | 0.00 |
| $\phi (Pop + Season)$ | 7 | 6827.97 | 13.84 | 0.00 |
| $\phi (Pop * Time)$ | 72 | 6845.92 | 31.80 | 0.00 |
| Only marble trout |  |  |  |  |
| $\phi (Pop + Time)$ | 26 | 6198.53 | 0.00 | 0.97 |
| $\phi (Pop)$ | 25 | 6207.06 | 8.53 | 0.01 |
| $\phi (Pop * Season)$ | 24 | 6207.89 | 9.36 | 0.00 |
| $\phi (Pop + Season)$ | 38 | 6208.59 | 10.06 | 0.00 |
| $\phi (Pop * Time)$ | 37 | 6223.68 | 25.15 | 0.00 |
| Only marble trout females with also spawning as predictor |  |  |  |  |
| $\phi (Pop + Time + Spawn\_temp)$ | 26 | 4558.48 | 0.00 | 0.47 |
| $\phi (Pop + Time)$ | 25 | 4558.59 | 0.11 | 0.45 |
| $\phi (Pop + Spawn\_temp)$ | 24 | 4564.64 | 6.16 | 0.02 |
| $\phi (Pop)$ | 38 | 4565.24 | 6.76 | 0.02 |
| $\phi (Pop + Season + Spawn\_temp)$ | 37 | 4566.12 | 7.64 | 0.01 |

We were able to confidently assign to parent-pairs or single parents 36% of genotyped individuals for UIdri_MT, 45% for LIdri_MT, and 25% for LIdri_RT (Fig. EMS2). Individuals continued to spawn throughout their lifetime (Fig. EMS3). The proportion of younger parents was higher in LIdri_MT than in UIdri_MT (χ^2^ = 4.98, *p* = 0.02), while no difference in proportion of younger parents was found between LIdri_MT and LIdri_RT (χ^2^ = 0.002, *p* = 0.96) (Fig. 4). Similar results were obtained when considering only 1+ as young parents.

**Fig. 4.**
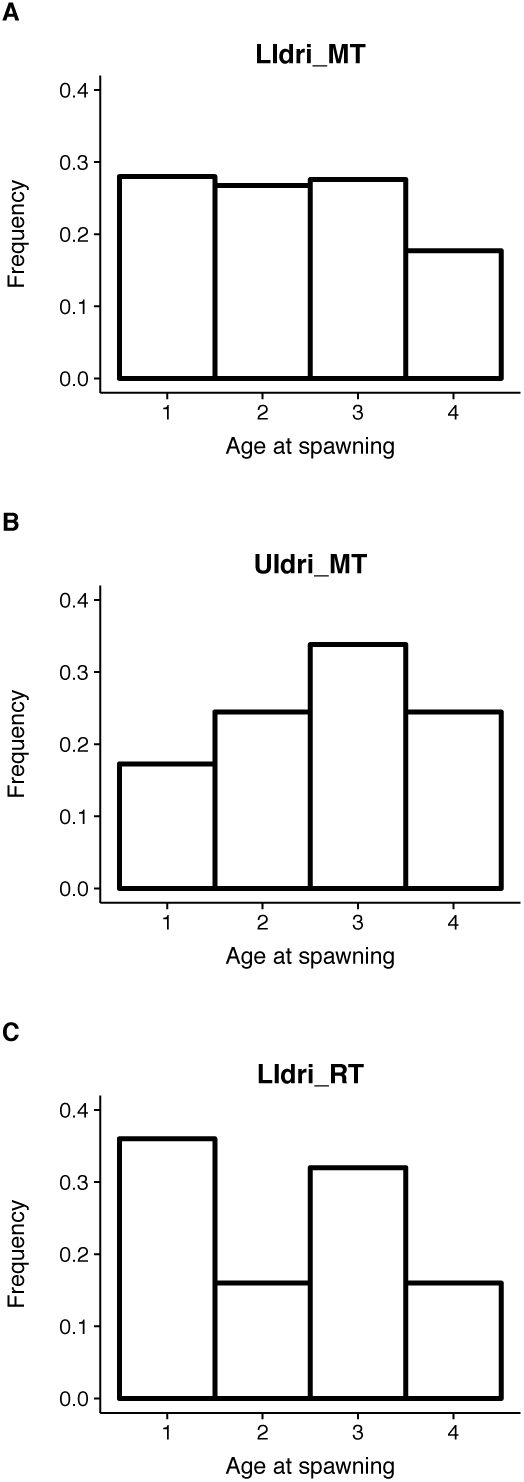
Age (up to 4+) distribution of parents in (a) LIdri_MT, (b) UIdri_MT, and (c) LIdri_RT.

## Discussion

The population dynamics of the two marble trout populations and of rainbow trout were overall synchronous. We found higher prevalence of younger parents, higher mortality, and lower population density in marble trout living in sympatry with rainbow trout than in marble trout living in allopatry. On the other hand, there were no differences in the average growth trajectories between the two marble trout populations. We discuss the observed variation in population dynamics, vital rates, and life histories between the two marble trout populations, the pieces of missing information that will improve our understanding of its proximate and ultimate causes, and how the observed variation is consistent with the predictions of life-history theory.

### Population dynamics and reproduction

In models of competition–colonization tradeoff, when two species occupying similar niches meet, the better competitor excludes the other (Cadotte 2007). On the other hand, evolutionary models of community assembly often assume that coexistence of competing species results from the operation of resource partitioning mechanisms (Macarthur and Levins 1967).

Musseau *et al.* (2016) found only a minor overlap between trophic niches of rainbow and marble trout in Lower Idrijca. They also found that marble trout in Upper and Lower Idrijca occupy a high trophic position (3.5 in Upper to 3.9 in Lower Idrijca), which confirms observations of piscivory and cannibalism in other marble trout populations. Marble trout in Lower Idrijca showed higher δ^13^C than marble trout in Upper Idrijca, which is likely to be caused by the inclusion of rainbow trout as additional prey in the diet of marble trout (Musseau et al. 2016). Our hypothesis is that predatory pressure on rainbow trout juveniles by marble trout older than 1+ caused both the low density of rainbow trout older than 0+ and the lack of cross-correlation between recruitment and density of older fish in rainbow trout. DNA-based faecal dietary analysis (Valentini and Al 2009) will help us test this hypothesis.

Overall, the population dynamics of marble trout in Upper and Lower Idrijca and of marble and rainbow trout in Lower Idrijca were synchronous, which points to common environmental determinants of temporal variation in vital rates (“Moran effect”). Vincenzi *et al.* (2016b) showed that variation in water temperature alone - over sampling occasions or early in life - did not explain much variation in either survival or growth in marble trout living in Western Slovenian streams, including the Idrijca populations. In addition, Vincenzi *et al.* (2016b) found strong cohort effects on survival and growth in almost all marble trout populations living in Western Slovenia, which suggest that a combination of environmental factors (e.g. food, water flow, temperature) experienced early in life is driving a large part of the variation in vital rates in marble and rainbow trout living in Idrijca. An analysis of spatio-temporal patterns of fluctuation of 57 brown trout populations widespread across France found that the degree of synchrony among sites for the 0 + fish was significantly related to the degree of hydrological synchrony (Cattaneo et al. 2003). Future measurement of water flows in Upper and Lower Idrijca with probes or meters and estimation of temporal variation of food-web components would provide a clearer picture of the environmental determinants of synchronous population dynamics.

At each sampling occasion, density of juveniles in marble trout in allopatry was much lower than that of marble trout in sympatry. The lower density of juveniles in allopatry was due to reproduction occurring mostly in a small tributary of Upper Idrijca located upstream of the two monitored stream stretches; the time passing between emergence of new-borns and sampling in September was not long enough to allow for the arrival of 0+ in the monitored stream stretches. It follows that estimated recruitment was only a small fraction of real recruitment.

### Survival

One of the central tenants of life-history theory is the expected trade-off between survival and reproduction. Vincenzi *et al.* (2016b) hypothesized that lower average survival in marble trout populations exhibiting faster individual growth was caused by the cost of reproduction: faster growth early in life can lead to younger age at sexual maturity when gonads development is size dependent (Alm 1959; Craig 1985; Jonsson et al. 2013) and then cause higher mortality of spawners due to energetic limitations (Berg et al. 1998). In marble trout living in Upper and Lower Idrijca, we did not find evidence of lower survival probabilities for female parents in the sampling interval (September to June) following spawning. However, most marble trout females spawn each year after becoming sexually mature and the cost of reproduction should not be limited to only those females that reproduced successfully. Thus, even if present, higher costs of reproduction in marble trout living in Lower Idrijca may be difficult to detect using parents as a proxy for spawners. Another hypothesis to explain lower survival of marble trout in sympatry with rainbow trout is more intense competition for hiding places; although the total density of fish in Upper and Lower Idrijca is similar, rainbow trout are bigger than marble trout and this may be due to their more aggressive and/or territorial behavior, with negative effects on marble trout survival rates.

### Age at spawning, growth, and life histories

We found that marble trout in sympatry had a higher proportion of young parents than marble trout living in allopatry, and the same proportion of young parents of sympatric rainbow trout. Along with lower survival, higher proportion of young parents points to faster life histories in sympatric than in allopatric marble trout.

It has often been found that fast life histories increase invasion success (Ruesink 2005) and that colonists adopt life histories that are faster than those of the source population (i.e. pioneer life histories), since younger age at reproduction and higher reproductive effort may facilitate rapid population growth and spatial expansion when the invading population is small (after the introduction) and throughout the invasion process (establishment and spread) (Burton et al. 2010; Kurz et al. 2016). Similarly, fast life histories in the invaded population may help avoid the invasion of the alien species or limit their abundance. For instance, Jones & Closs (2015) found that native galaxiid species (small freshwater fish living in the Southern Hemisphere) in New Zealand co-occurred with alien salmonids only when possessing fast life histories. In contrast, galaxiid species with slow life histories were excluded from salmonid-invaded reaches even when salmonid densities were low.

Theoretical and empirical studies have found that fitness is highly sensitive to variation in age at first reproduction (Fay et al. 2016). Age at reproduction varies greatly within- and among populations of the same species, and life-history theory predicts a trade-off between growth and reproduction (Stearns 1992; Roff 2007): organisms with a slower pace of life allocate more energy to growth throughout their life and reproduce at older ages, since fecundity and other reproductive traits, such as the ability to secure mates, often scale with size.

However, growth of fish results from the interaction of resource acquisition, resource availability, and resource allocation (Vincenzi et al. 2016a), along with other factors such as temperature, water flow, and length of growing season (Crozier et al. 2010). According to life-history theory, when there is minimum size for gonads development, organisms adopting faster life histories should grow faster and then plateau in size after sexual maturity due to allocation of energy to reproduction at the expense of growth. However, despite the presence of a competitor and the shift in diet, marble trout living in sympatry with rainbow trout did not grow differently than marble trout in allopatry, which shows that the acceleration of reproduction may occur without variation in growth.

Phenotypic shifts over contemporary time scales can result from phenotypic plasticity or selection acting on standing genetic variation, and they are most likely a result of a combination of these factors (Hendry and Gonzalez 2008). Reproductive traits are heritable (Carlson and Seamons 2008), and can thus respond adaptively to changes in the fitness landscape. In marble trout living in streams affected by flash floods causing massive mortalities, Vincenzi *et al.* (2017) found that fish born after the flash floods had younger mean age at reproduction than fish born before the flash flood. Vincenzi *et al.* (2017) hypothesized that younger at reproduction after the flash floods was due to a combination of faster growth due to lower population density and fewer older fish competing for mates. In the case of Idrijca, the age-structure of marble trout in Lower and Upper Idrijca and their fish densities are similar, and average growth trajectories are basically the same; thus other adaptive or plastic processes causing a higher prevalence of younger parents in marble trout living in Lower Idrijca may be at work. Currently on-going common garden experiments will help elucidate the proximal causes of faster life histories in marble trout living in Lower Idrijca.

## Acknowledgements

We thank the employees and members of the Tolmin Angling Association (Slovenia) for carrying out fieldwork since 1993. This study has been funded by MAVA Foundation.

## Authors’ Contributions

SV, AJC, and JCG conceived the ideas and designed methodology; AJC and DJ collected the data; SV and EC analyzed the data; SV, JCG, and AJC led the writing of the manuscript. All authors contributed critically to the drafts and gave final approval for publication.

## Data accessibiity

Genotype data: <u>https://dx.doi.org/10.6084/m9.figshare.4244252.v2</u>

Tag-recapture data: <u>https://dx.doi.org/10.6084/m9.figshare.4244249.v2</u>

